# Upregulation of voltage-gated potassium channels in peripheral blood mononuclear cells: a new approach to multiple sclerosis treatment

**DOI:** 10.1101/2025.02.03.636276

**Authors:** Marta Iglesias Martínez-Almeida, Ana Campos Ríos, Luís Freiría Martínez, Tania Rivera Baltanás, Daniela Rodrigues Amorim, César Manuel Sánchez Franco, Ainhoa Rodríguez Tébar, Mercedes Peleteiro, María Comis Tuche, Inés González Suárez, José Manuel Olivares, José Antonio Lamas, Carlos Spuch

## Abstract

Multiple sclerosis (MS) involves immune dysregulation and abnormal ion channel function. This study investigated the expression and function of voltage-gated potassium (Kv) channel isoforms (Kv1.1, Kv1.2, Kv1.3, Kv1.6, Kv4.2, Kv4.3, Kv7.2) in peripheral blood mononuclear cells (PBMCs) and lymphocytes from MS patients (remittent-recurrent) compared to controls. We found an upregulation of in six out of seven Kv isoforms in PBMCs from MS patients, with sex-specific differences observed (female showing a more pronounced upregulation of specific isoforms). Electrophysiological analysis of CD3+ T lymphocytes showed no significant differences, while CD19+ B lymphocytes displayed a notable decrease in outward current density. These findings suggest potential variations in Kv channel function across immune cell types and highlight the need for further exploration of their roles in MS pathogenesis, particularly within B lymphocytes and considering sex-based considerations.

## 1 Introduction

Multiple sclerosis (MS) is a chronic, autoimmune disease affecting millions worldwide, characterised by demyelination and neurodegeneration in the central nervous system (CNS) (Prinz and Priller, 2017). Despite significant research efforts, the exact aetiology and pathophysiology of MS remain inadequately understood. However, increasing evidence suggests the involvement of voltage-gated potassium (Kv) channels, particularly Kv1.3, in MS progression (Wang et al., 2020; Lioudyno et al., 2021).

Kv channels are crucial regulators of membrane potential and excitability in various cell types, including immune cells (Fellerhoff-Losch et al., 2016), in which modifications to either Kv channel expression or permeation can impact numerous physiological and functional characteristics such as regulating membrane potential, modulation of adhesion molecules activation mechanism along with regulating calcium influx, cytokine production, clonal expansion and cell death (Liu et al., 2002; Fellerhoff-Losch et al., 2016). Studies have shown upregulated Kv1.3 expression in T lymphocytes from MS patients compared to healthy controls, potentially contributing to disease pathogenesis (Wang et al., 2020). Additionally, polymorphisms in the Kv1.3 gene have been linked to faster MS progression (Lioudyno et al., 2021).

Given their established role in MS pathology, Kv channels present promising targets for therapeutic development. This study investigates the expression of various Kv channels (Kv1.1, Kv1.2, Kv1.3, Kv1.6, Kv4.2, Kv4.3, and Kv7.2) in PBMCs through Western blot analysis and Kv1.3 functional characteristics in peripheral blood T (CD3+) and B (CD19+) lymphocytes through patch-clamp electrophysiological recordings to assess potential differences between MS patients and controls.

## 2 Material and methods

### 2.1 Subjects and samples

Venous blood samples from 91 participants (39 patients diagnosed with RRMS and 40 healthy controls by the Western blotting; 7 patients diagnosed with MS and 5 healthy controls in electrophysiological recordings) were collected in vacuum tubes containing dipotassium ethylenediaminetetraacetic acid (K_2_EDTA) between 7:00 and 10:30 hours. The recruitment period lasted 1 year, and samples were obtained from the MS Unit at Álvaro Cunqueiro Hospital in Vigo, Spain. The RRMS diagnosis was achieved by neurologists based on a combination of clinical analysis, image probes, and cerebrospinal fluid analysis. The study’s inclusion criteria were patients with RRMS aged 18 years or older who provided a signed informed consent compliant with the guidelines of the Helsinki Declaration and approved by the ethics committee (Galician Network of Research Ethics Committees). Subjects who presented other neurological or psychiatric disorders were excluded from the study. Pregnant or breastfeeding women were also excluded. The control group was selected based on the frequency matching method (cases and controls have similar distributions of matching variables). Patients’ cohorts were matched by age and sex with a control group, which was composed of healthy volunteers aged 18 or older with no psychiatric or neurological disease and was also required to provide a signed informed consent. The MS Unit at neurology service used the standardised Expanded Disability Status Scale (EDSS) through a comprehensive physical assessment of muscle tone, strength balance, gait, coordination, vision, and cognitive function to measure the impact of the disease on the patient. The clinical and demographic characteristics of the study participants are shown in **Table 1**.

**Table 1.** Demographic and clinical data. Median and IQR. Legend: ^a^Mann-Whitney U test p-value; ^b^Fisher’s exact test p-value; ^c^years; ^d^NLR obtained from Forget et al., 2017. IQR: interquartile range. EDSS punctuation is between 0 and 10, where 0 is symptom absence and 10 death because of MS. Statistical significance: P≤0.05*, P < 0.01**; P <0.001***.

| Parameter | MS | Controls | P value |
| --- | --- | --- | --- |
| **Total, sample size (N)** | 46 | 45 |  |
| **Age (median and IQR) <sup>c</sup>** | 45 (42 – 53) | 50 (28 – 69) | 0.7979<sup>a</sup> |
| **Sex (M/F)** | 14/25 | 23/17 | 0.6466<sup>b</sup> |
| **Age of illness onset (median and IQR) <sup>c</sup>** | 32 (28 – 41.5) | - |  |
| **Duration of illness (median and IQR) <sup>c</sup>** | 10 (6.5 – 16.5) | - |  |
| **NLR (median and IQR)** | 2.71 (2.05 – 4.84) | 1.66 (1.26 – 2.16)<sup>d</sup> | < 0.0001<sup>a ***</sup> |
| **EDSS** | 2.5 (2 – 5.75) | - |  |

### 2.2 Sample preparation

Venous blood is collected into two vacuum tubes containing K_2_EDTA (BD vacutainer; Becton, Dickinson & Co., USA). The blood is transferred to a centrifuge tube containing lymphocyte isolation solution (Rafer, Zaragoza, Spain) and centrifuged at 2000 rpm for 35 min to separate peripheral blood mononuclear cells (PBMCs), which were subsequently washed 2 times with phosphate buffered saline (PBS) at 1500 rpm for 5 minutes. The pelleted PBMCs are resuspended in PBS and divided into three aliquots, one is then stored at −80 °C. One fresh aliquot of PBMCs is used to isolate CD3+ and CD19+ lymphocyte subpopulations using magnetic beads and then are used for patch-clamp and immunofluorescence. For immunofluorescence, lymphocytes are fixed with paraformaldehyde (PFA) 4 % 5 min and then centrifuged 10 min at 1500 rpm. Afterwards the PBMCs are once again washed 2 times with PBS at 1500 rpm for 5 min and then used for immunofluorescence.

### 2.3 CD3+ and CD19+ isolation

Commercial magnetic bead kits were used to isolate CD3+ lymphocytes (Dynabeads Untouched Human T-cells; 11344D; Invitrogen, Thermo Fisher Scientific, Waltham, MA, USA) and CD19+ lymphocytes (Dynabeads Untouched Human B-cells; 11351D; Invitrogen, Thermo Fisher Scientific, Waltham, MA, USA) from PBMCs. The procedures were performed following the manufacturer’s instructions. Isolated lymphocytes are divided into two aliquots to perform immunofluorescence and electrophysiological recordings.

### 2.4 Isolation validation by flow cytometry

The isolation of cell subtypes using the magnetic beads was verified by means of flow cytometry. After isolation, the cells were centrifuged at 1200 rpm for 5 min at room temperature, the supernatant was removed, and the cells were resuspended in phosphate buffer saline (PBS) containing 0.5 % bovine serum albumin (BSA). 500,000 cells were distributed per vial and incubated with 10 µl of receptors of constant fraction (FcR) blocking reagent (FCR-2ML; Immunostep S.L.) for 10 min. CD3-FITC (MA1-10178; Thermo Fisher Scientific, Waltham, MA, USA) and CD19-ECD (A07770; Beckman Coulter, Inc.) antibodies were then added and incubated for 30 min at 4 °C. The cells are then centrifuged at 1200 rpm for 5 min at room temperature, the supernatant is removed, and the cells are resuspended in 100 µl of PBS containing 0.5 % BSA. The cell volume was passed through a 0.4 µm filter and cytometry was performed in CytoFLEX S (Beckman Coulter, Inc.) flow cytometer at the Flow Cytometry facilities from the CINBIO, University of Vigo. Acquisition of flow cytometry data involved the recording of at least 10,000 events per sample and subsequent analysis, both using CytExpert Software version 2.3.1.22 (Beckman Coulter, Inc.). Following the initial gating based on forward and side scatter plots to isolate lymphocytes, singlets were selected using forward scatter area vs. forward scatter height criteria. Recorded data from cell populations were expressed as percentages of total cells counted. **Supplementary figure 1** and **Supplementary figure 2** provide details on gating strategies employed for identifying different subsets within each panel studied.

### 2.5 Immunofluorescence microscopy

Depart from fixed PBMCs, the supernatant is discarded, and cells are resuspended in Envision Flex Antibody diluent (Dako, Denmark) at 4 °C and then centrifuged at 1500 rpm 10 min at 4 °C. Supernatant is discarded, cells are resuspended in 5% BSA and got into rotation 30 min at 4 °C. After blocking cells are centrifuged 10 min 1500 rpm at 4 °C. Supernatant is discarded and cells are resuspended in 0.1 % Triton-×100 and overnight incubation with primary antibodies (Table 2) in a rotating mixer.

**Table 2.** Primary antibodies used in immunofluorescence procedures. Kv1.1, voltage-gated potassium channel subfamily A member 1; Kv1.2, voltage-gated potassium channel subfamily A member 2; Kv1.3, voltage-gated potassium channel subfamily A member 3; Kv1.6, voltage-gated potassium channel subfamily A member 6; Kv4.2, voltage-gated potassium channel subfamily D member 2; Kv4.3, voltage-gated potassium channel subfamily D member 3; Kv7.2, voltage-gated potassium channel subfamily KQT member 2.

| Antibody | Origin | Type of antibody | Trading house | Concentration |
| --- | --- | --- | --- | --- |
| **Anti-Kv1.1** | Rabbit | Polyclonal | Abcam plc | 1:250 |
| **Anti-Kv1.2** | Mouse | Monoclonal | Abcam plc | 2.5 µg/ml |
| **Anti-Kv1.3** | Rabbit | Polyclonal | Abcam plc | 1:250 |
| **Anti-Kv1.6** | Rabbit | Polyclonal | Abcam plc | 0.5 µg/ml |
| **Anti-Kv4.2** | Rabbit | Polyclonal | Abcam plc | 1:100 |
| **Anti-Kv4.3** | Rabbit | Polyclonal | Abcam plc | 1:100 |
| **Anti-Kv7.2** | Mouse | Monoclonal | Abcam plc | 5 µg/ml |
| **Anti-CD3** | Rabbit | Polyclonal | Abcam plc | 1:100 |
| **Anti-CD3 FITC** | Mouse | Monoclonal | Thermo Fisher Scientific | 1:100 |

Aliquot is again centrifuged 10 min 1500 rpm at 4 °C, supernatant is discarded, and cells resuspended in 1 % Tris-buffered saline (TBS; pH 7.5); centrifugation is repeated. Supernatant is discarded again, and cells are resuspended in 0.1 % Triton-×100 with secondary antibodies listed in Table 3.

**Table 3.**
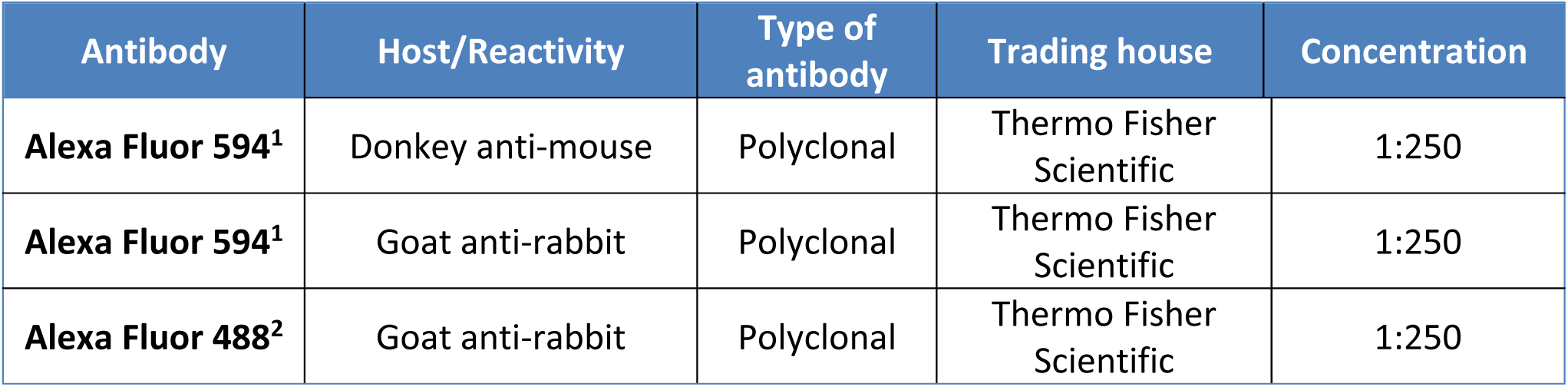
Secondary antibodies used in immunofluorescence. ^1^ Red immunofluorescence analysis (excitation peak at 490 nm; emission peak at 617 nm). ^2^ Green immunofluorescence analysis (excitation peak at 490 nm; emission peak at 525 nm).

Aliquots are incubated in rotation 2 hours at 4 °C, after this time they are centrifuged 10 min 1500 rpm at 4 °C. Centrifugation is repeated after resuspending cells in TBS 1%, supernatant is discarded and 4,6-diamidino-2-phenilindol (DAPI; Sigma; 1:1000) is added and incubated for 15 min. Aliquots are centrifuged 2× with 1 % TBS 10 min 1500 rpm at 4 °C. Finally, supernatant is discarded, and cells are pipetted in Superfrost® Plus Slides (Thermo Fisher Scientific, Waltham, MA, USA) with mounting medium Mowiol® (Sigma Aldrich, MO, USA) and covered with a coverslip. Samples were visualised with Leica Stellaris 8 100.0 × 3.5 OIL objective controlled by Leica LAS X software.

### 2.6 Western blot

Protein samples were mixed with 2× volume of Laemmli buffer 2× (Bio-Rad, California, USA) (950 µl of Laemmli buffer and 50 µl of β-mercaptoethanol) and lysis buffer PIK, and boiled at 95 °C for 5 min. Fraction samples were loaded in 8 % Bis-Tris polyacrylamide gels, and an electrophoresis was performed in a Power Pac ^TM^ universal power supply (Bio-Rad, California, USA) at 80 V for 30 min, and at 120 V for another 90 min. The proteins were immediately transferred to polyvinylidene difluoride membranes (Immun-Blot ® polyvinylidene fluoride (PVDF) membrane, Bio-Rad, California, USA) contained in a PowerPac^TM^ universal power supply (Bio-Rad, California, USA) set at 0.25 A for 60 min (for two gels). The membranes were blocked with 5% milk powder in a tris-buffered saline solution with Tween (TBST) for 20 min and washed three times with TBST solution. The membranes were incubated overnight stirring at 4 °C with a primary antibody listed in Table 4.

**Table 4.**
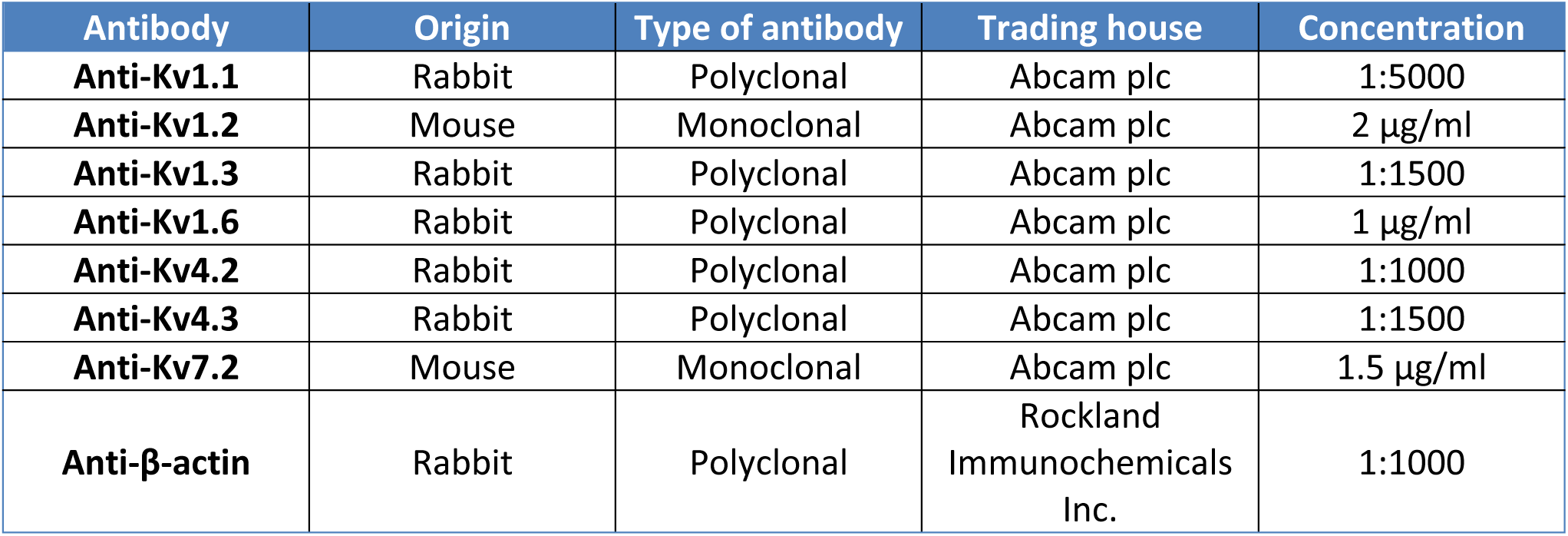
Primary antibodies used in Western blot.

| Antibody | Origin | Type of antibody | Trading house | Concentration |
| --- | --- | --- | --- | --- |
| **Anti-Kv1.1** | Rabbit | Polyclonal | Abcam plc | 1:5000 |
| **Anti-Kv1.2** | Mouse | Monoclonal | Abcam plc | 2 µg/ml |
| **Anti-Kv1.3** | Rabbit | Polyclonal | Abcam plc | 1:1500 |
| **Anti-Kv1.6** | Rabbit | Polyclonal | Abcam plc | 1 µg/ml |
| **Anti-Kv4.2** | Rabbit | Polyclonal | Abcam plc | 1:1000 |
| **Anti-Kv4.3** | Rabbit | Polyclonal | Abcam plc | 1:1500 |
| **Anti-Kv7.2** | Mouse | Monoclonal | Abcam plc | 1.5 µg/ml |
| **Anti-β-actin** | Rabbit | Polyclonal | Rockland Immunochemicals Inc. | 1:1000 |

After washing the membranes three times with TBST, they were incubated with appropriate secondary antibody 1:10,000 (GE Healthcare Life Sciences, UK) stirring during 60 min. The membranes were then washed again twice with TBST and once with TBS. The ChemiDoc XRS+ system (Bio-Rad, California, USA) was then used to analyse the chemiluminescence of the membranes with the ECL^TM^ Prime Western Blotting System (Sigma-Aldrich, St. Louise, USA). Image Lab 6.0 software (Bio-Rad, California, USA) was used to analyse the blot images acquired, and a densitometric ban quantification was performed by means of the Image J 1.5K software (National Institutes of Health, USA). Western blot normalisation was performed individually using β-actin for PBMCs, converting results in percentage expressed in relative units.

### 2.7 Electrophysiological recordings

Whole-cell patch-clamp experiments were performed in the Laboratory of Neuroscience facilities from the CINBIO (University of Vigo) 1 hour after CD3+ and CD19+ were freshly isolated and plated in 35 mm wells previously coated with laminin 10 µg/ml. The standard extracellular solution contained 135 mM NaCl, 5 mM KCl, 1 mM CaCl_2_, 1 mM MgCl_2_ and 10 mM HEPES (pH 7.4 and osmolality 270-280 mOsm/kg) and the intracellular solution, 100 mM KCl, 40 mM KF, 1 mM CaCl_2_, 1 mM MgCl_2_, 10 mM HEPES and 10 mM EGTA (pH 7.4 and osmolality 290 mOsm/kg) as previously described by Markakis et al., 2021. Patch pipettes were fabricated from borosilicate capillaries (Harvard Aparattus, Massachusetts, USA) using a P-1000 horizontal puller (Sutter Instrument, California, USA), with resistances between 3 and 6 MΩ. All recordings were performed at room temperature (20-24 °C) and currents were low pass filtered at 2 kHz. To study Kv currents in CD3+ and CD19+ cells in culture, voltage ramps from −100 to 100 mV (150 ms) were recorded and voltage jumps from −80 to 40 mV (1000 ms, 15 mV increments) were performed to construct conductance-to-voltage curves (fitted to Boltzman function), where conductance was calculated based on the recorded current and using the Ohm’s Law. In both cases, the membrane potential was held at −80 mV. Peak currents were measured at 40 mV. Measurements were accomplished by using pClamp software (Molecular Devices, California, USA).

### 2.8 Statistical analysis

The GraphPad Prism 8 software (GraphPad Software Inc., San Diego, CA, USA) was used to manage the resulting data and to perform the statistical analysis from western blot protein quantification. The mean age of both cohorts was compared with the Mann-Whitney U test, and the differences between sex ratios were analysed with Fisher’s exact test. Each band of each blot was analysed individually and normalised based on the average of controls of each gel, whose results were converted in percentage and expressed in relative units. The Shapiro-Wilk test was used to determine whether the results confirmed the normality. As this was not the case, non-parametric tests were used: the Mann-Whitney test was used for the median and interquartile range (IQR) comparison between groups and the Spearman analysis for correlations. Two-way analysis of variance (Geisser Greenhouse correction) with Bonferroni *post hoc* correction was applied to assess multiple comparisons in electrophysiological experiments. Results are considered statistically significant when *P-*value ≤ 0.05.

## 3 Results

### 3.1 Demographic and clinical data analysis

A case-control study was performed to compare a cohort of patients with MS and a cohort of healthy subjects. The comparison of the demographic and clinical data of 39 patients with RRMS and 40 healthy subjects revealed no significant differences between groups regarding age (*p* = 0.52218), and sex (*p* = 0.0784) (Table 1). The neutrophil-lymphocyte ratio (NLR) is a parameter used to easily assess the systemic inflammation of a subject. Accordingly, the NLR was quantified in patients, and a standard value was used for the control cohort (Forget et al., 2017). The comparison revealed that there were no statistically significant differences between groups (*p* = 0.4511) (Table 1). Additionally, a correlation analysis was conducted to investigate the correlation between levels of Kv channels, and clinical variables, such as EDSS, age of illness onset, duration of illness, or NLR of the group of patients with MS.

### 3.2 CD3+ and CD19+ magnetic beads isolation validation

The CD3+ and CD19+ isolation was validated using flow cytometry to verify the purity of the cell subtype. The results are shown as percentages in Table 5.

**Table 5.** Isolation validation percentages by flow cytometry.

|  | CD3+ | CD19+ |
| --- | --- | --- |
| **Isolated CD3+ control** | 92,35 % | 0,01 % |
| **Isolated CD19+ control** | 0,09 % | 98,67 % |
| **Isolated CD3+ MS** | 94,51 % | 0,01 % |
| **Isolated CD19+ MS** | 0,15 % | 98,41 % |

### 3.3 Immunofluorescence microscopy

To investigate the presence and subcellular distribution of Kv channels in lymphocytes from healthy individuals and those suffering from MS, immunostaining was conducted. The staining procedure involved blue dyeing of the nucleus, triple-labelling T lymphocytes for each Kv with red coloration, green labelling CD3+ antigenic marker, and nuclear labelling; while B lymphocytes were double-labelled for Kv using red coloration, along with blue nuclear marking. However, due to complications during experimentation only one type of Kv (Kv1.1) could be stained in B lymphocytes. Previous information on the presence and distribution of these channels in lymphocytes was extracted from the Protein Atlas and Uniprot databases, where all voltage-gated potassium channels except Kv4.2 were found to be present in T and B lymphocytes.

In T lymphocytes, all voltage-gated potassium channels were found, including Kv4.2 (Figure 1). In B lymphocytes, Kv1.1 was found to be present (Figure 2) and although it was not possible to obtain fluorescence images for the remaining Kv, our previous studies suggest that Kv1.1, Kv1.3 and Kv4.2 are present in these cells, while the information in the Protein Atlas and Uniprot suggests that the remaining Kv studied are also present. The localization and distribution of proteins in the cells does not appear to differ between controls and MS patients cells.

**Figure 1.**
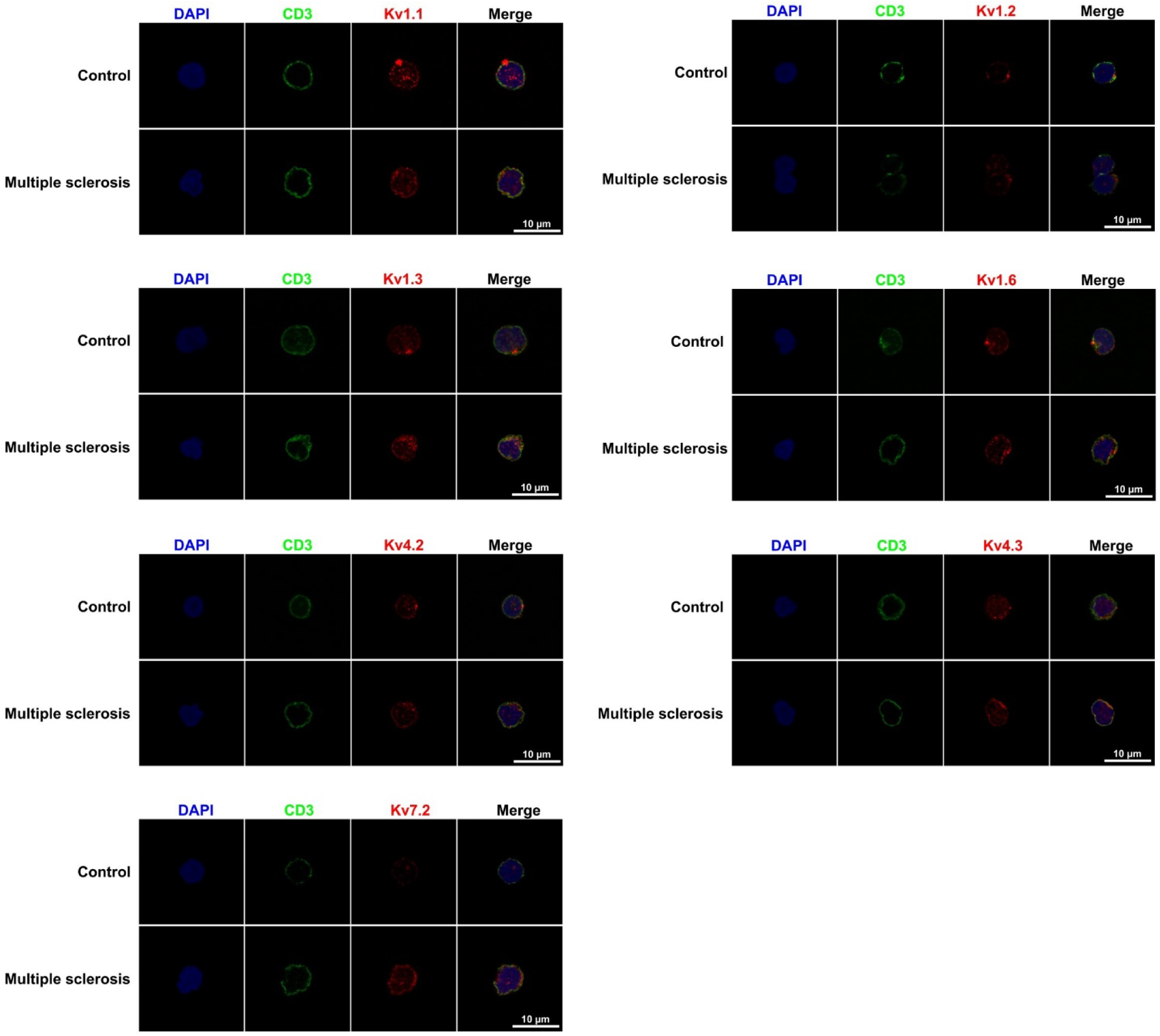
T lymphocyte immunostaining. Fluorescence microscopy images of blood T lymphocytes from healthy participants (control) and MS participants red colour labelled with Kv1.1, Kv1.2, Kv1.3, Kv1.6, Kv4.2, Kv4.3 and Kv7.2; and green colour labelled with CD3+. The white scale bar represents 10 µm.

**Figure 2.**
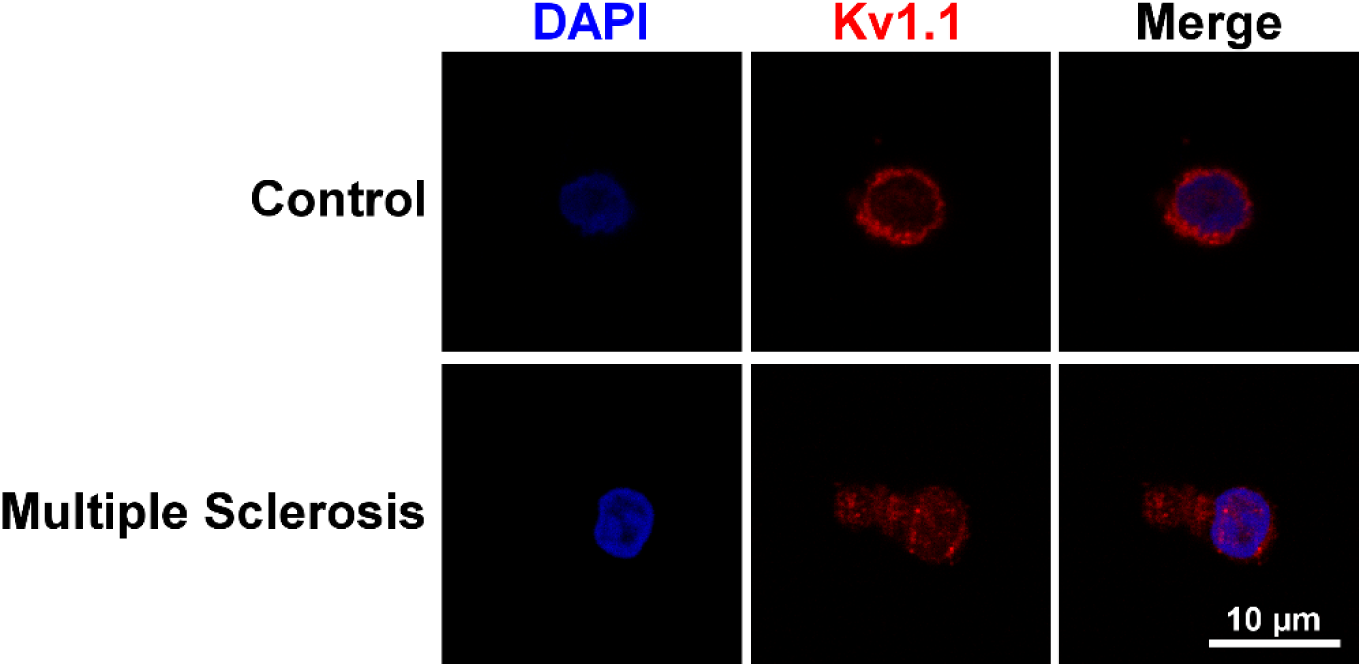
B lymphocyte immunostaining. Fluorescence microscopy images of blood B lymphocytes from healthy participants (control) and MS participants red colour labelled with Kv1.1. The white scale bar represents 10 µm.

### 3.4 Voltage-gated potassium channel quantification

A case-control study was conducted to investigate PBMCs of individuals diagnosed with MS and healthy participants through the application of western blot technique. The evaluation of PBMC levels involved determining their relative units, which were obtained by contrasting mean percentage values against controls in both groups. The statistical analysis of protein levels in PBMCs revealed significant higher levels in MS patients than in controls for Kv1.1 (*p* = 0.0002), Kv1.2 (*p* < 0.0001), Kv1.3 (*p* = 0.0003), Kv1.6 (*p* = 0.0005), Kv4.3 (*p* = 0.0047) and Kv7.2 (*p* = 0.0027) (Figure 3). Additional statistical values are summarised in Supplementary Table 1.

**Figure 3.**
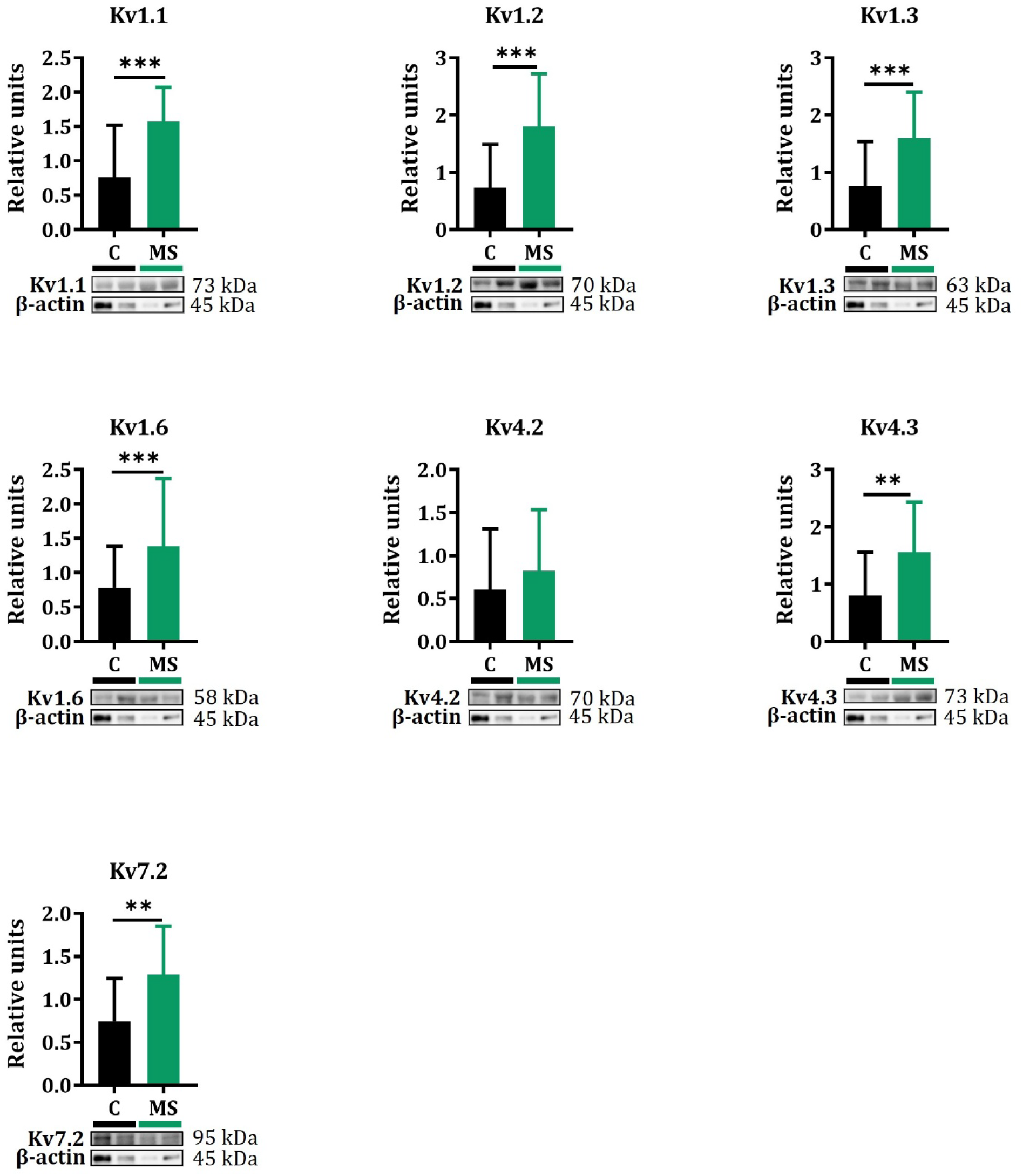
Protein expression (median and interquartile range) and representative western blots results in PBMCs from patients with MS (n = 40) and healthy controls (n = 40). Kv levels were measured by western blot and normalised with the housekeeping protein β-actin. Relative units refer to the mean of control values after normalisation. Statistically significant according to Mann-Whitney U test: P ≤ 0.05*, P < 0.01**; P <0.001***.The results show statistical significance differences between MS and C for Kv1.1 (p = 0.0002), Kv1.2 (p < 0.0001), Kv1.3 (p = 0.0003), Kv1.6 (p = 0.0005), Kv4.2 (p = 0.2674), Kv4.3 (p = 0.0047) and Kv7.2 (p = 0.0027).

Our study analysed the potential influence of sex on protein expression in PBMCs. We conducted analyses to contrast healthy males and those with MS, as well as healthy females and those diagnosed with MS (Figure 4). Additionally, we compared protein levels across sex within both control groups and MS cohorts. A significant overexpression of Kv1.1 was found in MS men (*p* = 0.0114) and women (*p* = 0.0230) compared to healthy participants; for Kv1.2 significant overexpression was found in MS women compared to controls (*p* = 0.0003); for Kv1.3 significant higher levels were found in MS men (*p* = 0.0152) and women (*p* = 0.0303) compared to healthy subjects; for Kv1.6 MS men showed higher levels than control men (*p* = 0.0282); for Kv4.3 MS women show an overexpression compared to healthy participants (*p* = 0.0225), and MS women also show significant higher levels than MS men (*p* = 0.0465); and in Kv7.2 there was found significant higher levels in MS women compared to controls (*p* = 0.0228) and MS women also showed overexpression compared to MS men (*p* = 0.0345). Additional statistical data can be found in Supplementary Table 2.

**Figure 4.**
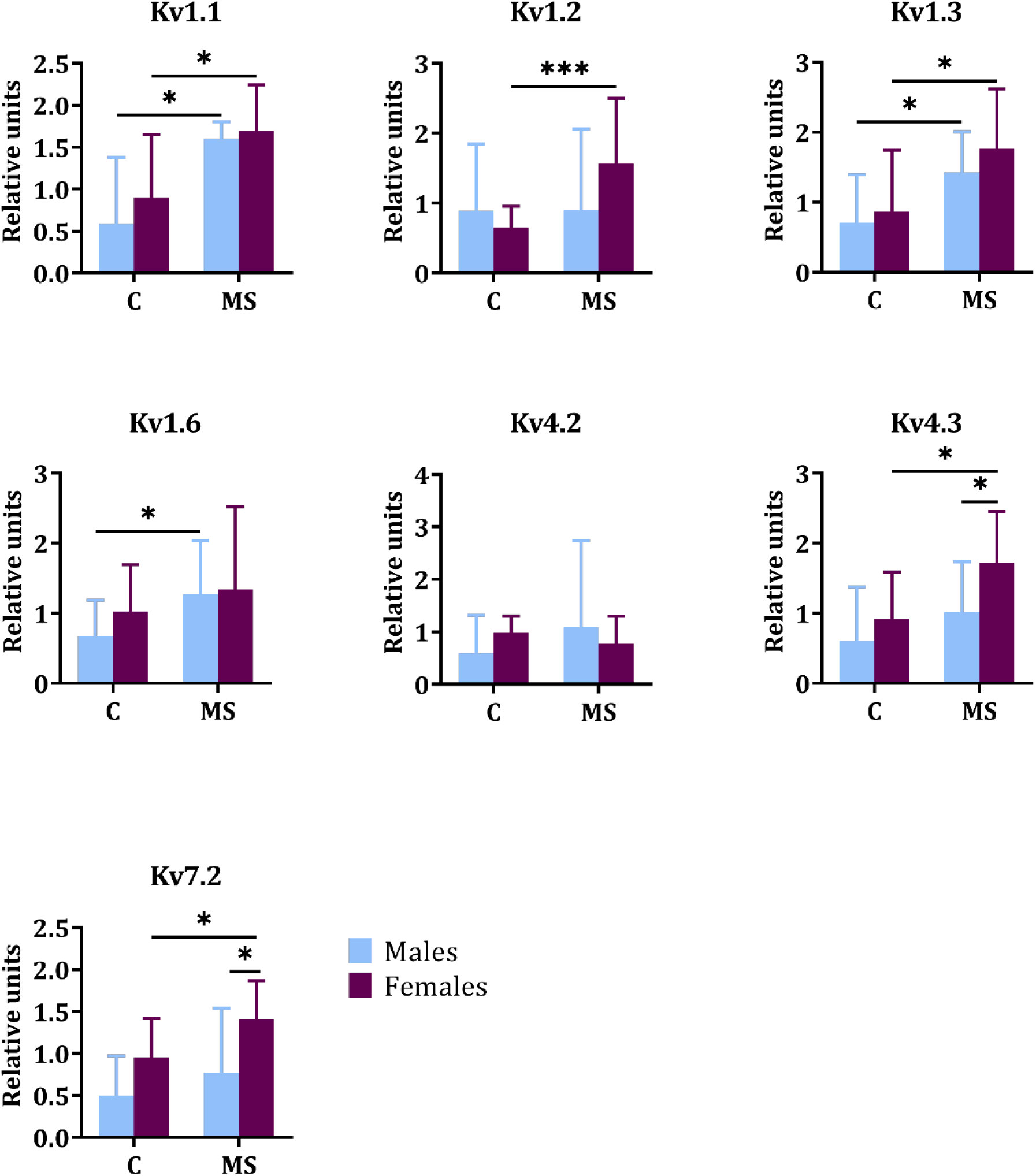
Sex comparison of PBMCs protein expression between SZ and. **C.** Plots represented by median and interquartile range. Statistically significant according to Mann-Whitney U test: P ≤ 0.05*, P < 0.01**; P <0.001***.

No correlations were found with clinical variables (Supplementary Table 3). No significant differences were found in protein levels between different MS types, such as relapsing remitting (RRMS), secondary progressive, and primary progressive. Either way, there were not found significant differences in protein expression between the different treatments.

### 3.5 Electrophysiological recordings

To investigate whether differences found in Kv1.3 protein expression between MS patients and healthy controls had a direct impact on Kv1.3 currents, we performed whole-cell patch clamp experiments in freshly isolated T and B lymphocytes, respectively.

The presence of cumulative inactivation (see Figure 5A) is a distinctive property of Kv1.3 currents, so the currents observed by recording outward currents in response to −100 to 100 mV ramps (150 ms) and to −80 to 40 mV jumps (15 mV increments, 1000 ms) might be identified as Kv1.3. Regarding T lymphocytes (Figure 5A), no differences were found between recorded ramps from both groups. Peak current amplitudes were also observed to be similar between healthy controls and MS individuals (Figure 5B). Conductance-to-voltage relation showed that these currents have similar biophysical characteristics and no differences in conductance between groups (Figure 5C), suggesting that the upregulation in the channel functional expression might not affect their electrophysiology.

**Figure 5.**
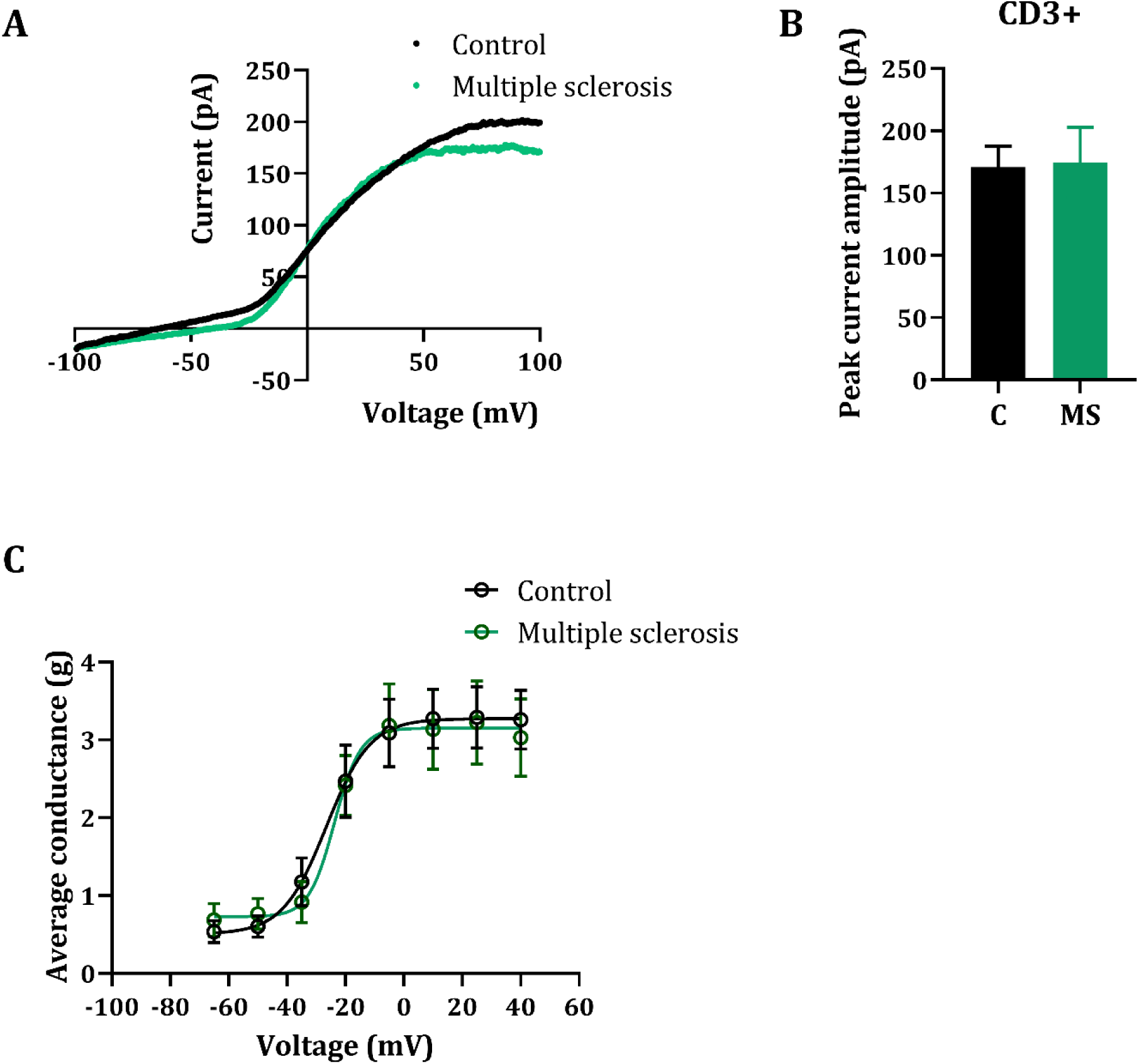
Patch-clamp data from T lymphocytes in MS patients compared to healthy individuals. (A) Current average response to a −100 to 100 mV ramp (150 ms) at a holding potential of −80 mV in both groups. No differences between ramps were found (two-way analysis of variance with multiple comparisons). (B) Peak current amplitudes are also similar between healthy controls (n = 5 individuals, N = 10 cells) and MS patients (n = 7 individuals, N = 9 cells). Data shown are average values ± S.E. (C) Peak amplitudes in response to 1000 ms jumps from −80 to 40 mV (15 mV increments) were converted to conductance-to-voltage curve by fitting to Boltzmann function (conductance calculated with Ohm’s law). Note that average Kv conductance is similar in both groups. The average peak conductance at 40 mV was 3.25 ± 0.38 nS in healthy controls and 3.02 ± 0.5 in MS patients (Mann-Whitney test comparison, p = 0.72).

Nevertheless, when comparing ramps from B lymphocytes (Figure 6A), we observed that MS patients presented a decrease in the outward current, starting at a membrane potential of 30 mV (*p* = 0.0236) and lower peak current amplitudes (*p* = 0.0102) (Figure 6B). Surprisingly, conductance-to-voltage fitted curves were similar in both groups (Figure 6C). Despite the predicted upregulation in the western blot analysis, these data show a clear reduction in outward currents through Kv channels in B lymphocytes, which suggest that these channels might present different dysfunctions in different PBMC types.

**Figure 6.**
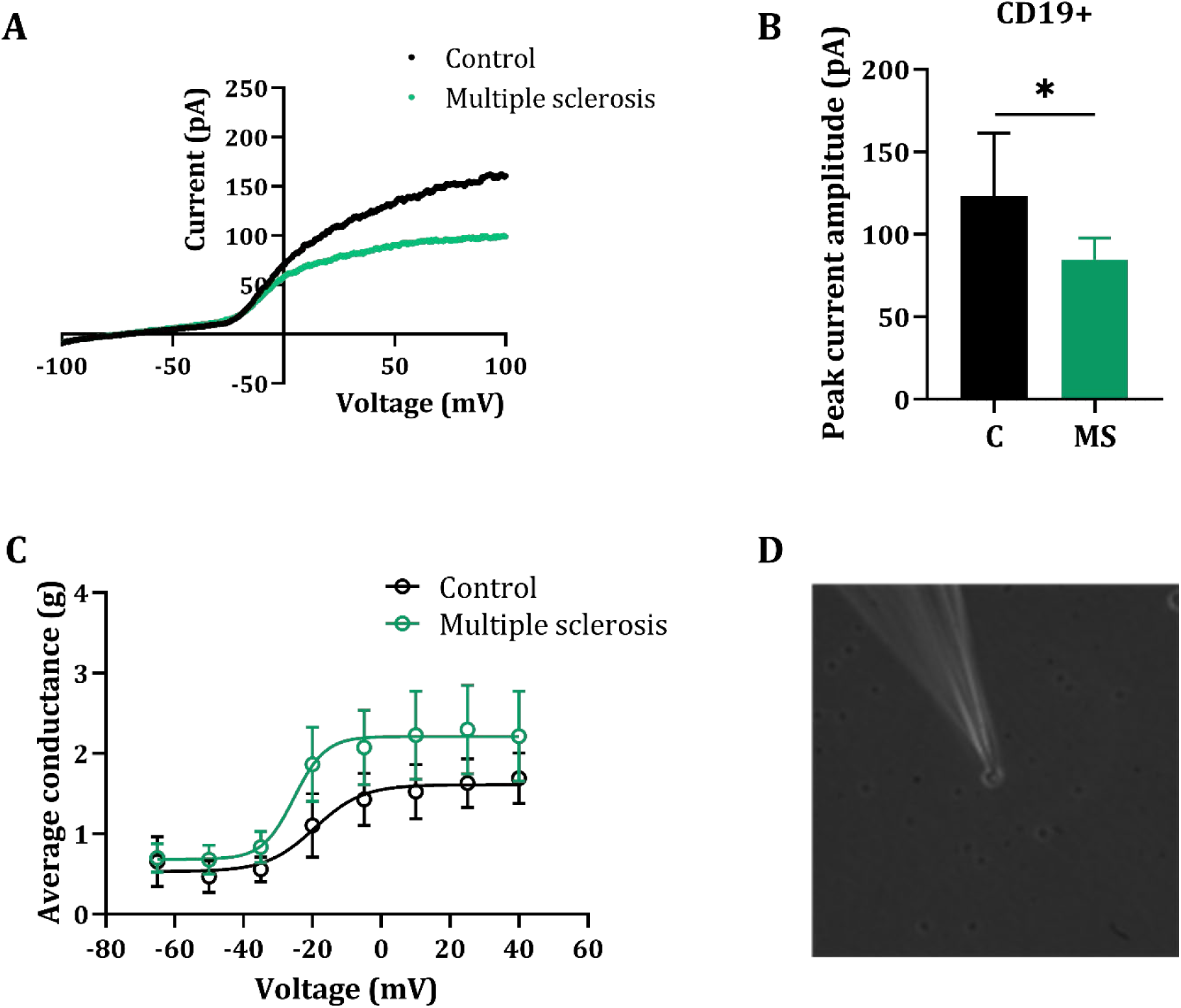
Patch-clamp data from B lymphocytes in MS patients compared to healthy individuals. (A) Current average response to a −100 to 100 mV ramp (150 ms) at a holding potential of −80 mV in both groups. B lymphocytes present lower outward currents, starting from 30 mV membrane potential (p = 0.0236) (two-way analysis of variance with multiple comparisons). (B) Peak current amplitude is also decreased (p = 0.0102) in MS patients (n = 7 individuals, N = 7 cells) compared to healthy controls (n = 5 individuals, N = 5 cells). Data shown are average values ± S.E. (C) Peak amplitudes in response to 1000 ms jumps from −80 to 40 mV (15 mV increments) were converted to conductance-to-voltage curve by fitting to the Boltzmann function (conductance calculated with the Ohm’s law). Note that average Kv conductance is similar in both groups. The average peak conductance at 40 mV was 1.69 ± 0.31 nS in healthy controls and 2.21 ± 0.56 in MS patients (Mann-Whitney test comparison, p = 0.44). (D) CD19+ cell from a MS patient while being recorded.

## 4 Discussion

Abnormalities in ion channels, such as an altered expression or functionality, in the CNS and in the immune system, are one of the mechanisms of immune-mediated damage in MS (Schattling et al., 2014; Wang et al., 2020). Due to the limited understanding of the role that Kv channels play in the CNS and immune system in MS, our research aims to expand knowledge on this topic and, therefore, we investigated the expression of various Kv channel isoforms (Kv1.1, Kv1.2, Kv1.3, Kv1.6, Kv4.2, Kv4.3 and Kv7.2) in PBMCs from MS patients and healthy controls, as well as their distribution on CD3+ and CD19+ using fluorescence techniques.

Our findings revealed an upregulation of six out of seven studied Kv channel isoforms in PBMCs from MS patients compared to controls (Figure 3). This observation adds to the growing body of evidence suggesting that Kv channels, particularly Kv1.3, play a critical role in MS pathogenesis. Notably, this is the first study to report the overexpression of these Kv channel isoforms in PBMCs from MS patients. Impaired normal function of Kv channels, usually by overactivation or overexpression, has been demonstrated in chronic inflammatory diseases, such as MS (Teisseyre et al., 2022). Previous research has predominantly focused on the investigation of Kv1.3, a channel that is usually overexpressed in CNS and in T lymphocytes from patients with MS (Rangaraju et al., 2009; Markakis et al., 2021); our observations are consistent with this as we also detected Kv1.3 overexpression in PBMCs from MS patients (Figure 3). In individuals with MS, the blood brain barrier (BBB) is compromised resulting in the invasion of peripheral immune system cells, as myelin-reactive T lymphocytes and memory effector T lymphocytes, that exhibit an overexpression of Kv1.3 (Hu et al., 2013; Wang et al., 2020). Upon activation, lymphocytes release calcium ions that were stored in the endoplasmic reticulum into the cytosol. This action leads to a calcium influx through specific channels, causing membrane potential depolarization and activating Kv1.3, responsible for restoring the resting membrane potential by facilitating potassium efflux (Hu et al., 2013; Kazama et al., 2015; Lee et al., 2024). These coordinated mechanisms control subsequent immune system responses; therefore, the upregulation of Kv1.3 leads to an overactivation of intracellular calcium signalling that promotes cell activation, proliferation and cytokine production, thereby highlighting its potential involvement in inflammatory processes related to MS (Wang et al., 2020).

Moreover, this mechanism is observed in neurons as well, where heightened excitability and raised intracellular calcium levels result in impaired mitochondrial function and neuronal death (Bozic et al., 2018). In support for this theory lies evidence from various studies showing that inhibition or blocking Kv1.3 could reduce T lymphocyte response while keeping BBB functions intact and suppressing autoimmune neuroinflammation (Rangaraju et al., 2009; Zhao et al., 2020).

Our study also identified sex-specific differences in Kv channel expression, with females exhibiting a more pronounced upregulation of Kv1.1, Kv1.2, Kv1.3, Kv4.3, and Kv7.2 compared to control females; and men exhibiting an upregulation of Kv1.1, Kv1.3, and Kv1.6 compared to their control counterparts (Figure 4). Additionally, women with MS present an upregulation of Kv4.3 and Kv7.2 compared to men with MS (Figure 4). Progesterone and oestradiol are hormones that have been found to block Kv channels in various cell types in females. Kv channels in T lymphocytes are directly blocked by progesterone, which leads to membrane depolarization, inhibiting calcium signalling pathways and decreasing the production of cytokines. As a result, progesterone contributes towards immunosuppressive effects (Ehring et al., 1998). In MS, progesterone showed neuroprotective characteristics by reducing axonal damage and promoting oligodendrocyte myelin repair (Bridge et al., 2023). It has been hypothesised that such benefits could be achieved through the inhibition of Kv channels by these hormones. Testosterone, another crucial sex hormone, experiences significant reduction in individuals with MS regardless of sex. Research indicates that women and men with lower levels of testosterone present a higher number of lesions and cognitive dysfunction respectively (Bove et al., 2014; Bridge et al., 2023). Emerging evidence highlights the potential link between testosterone deprivation, impairment of Kv channel currents as well as modification in its expression patterns observed across diverse animal models and cell lines (Baldwin et al., 2023). These findings warrant further investigation into the potential influence of sex hormones, such as progesterone, oestradiol, and testosterone, on Kv channel expression in MS, considering their known modulatory effects in various cell types.

While electrophysiological studies on Kv channels have been conducted in the nervous system and experimental autoimmune encephalomyelitis models (Beeton et al., 2001; Gentile et al., 2020), research on immune cells in MS remains limited and may provide an insight into the functional role of specific channels, which is of special interest to the understanding of immune-related diseases as well as for developing potential therapeutic interventions. Our study addressed this gap by examining Kv currents in both CD3+ (Figure 5) and CD19+ (Figure 6) lymphocytes from relapsing remitting MS patients. Interestingly, contrary to our initial hypothesis based on Western blot data, significant differences were not observed in CD3+ T lymphocytes (Figure 5), while CD19+ B lymphocytes displayed a notable decrease in outward current density (Figure 6A) and significant lower peak amplitudes (Figure 6B). These findings suggest potential variations in Kv channel functions across different immune cell types and highlight the need for further exploration of their roles in MS pathogenesis. Upon reaching this point, further research using specific pharmacological blockers might be interesting to deepen in the function of particular Kv channels in the pathophysiology of MS.

The importance of Kv channels in T lymphocytes has already been established in MS (Zhang et al., 2016; Chisari et al., 2022). However, their function is not fully comprehended in B lymphocytes, despite being linked to cellular activation, proliferation, differentiation and cytokine production (Wang et al., 2012; Zhang et al., 2016). Furthermore, there is limited research on the role of Kv channels in MS disease. Interestingly, rituximab, a medication used for treating MS and lymphomas, has been shown to inhibit Kv1.3, leading to lymphoma cell death (Wang et al., 2012). While the specific impact of rituximab on Kv1.3 in MS remains to be elucidated, its effectiveness in reducing inflammation, relapse frequency, and CNS lesions suggests a potential link to B lymphocyte function (Chisari et al., 2022). This highlights the need for further investigation into the role of Kv channels in B lymphocytes within the context of MS.

The use of rituximab in treating lymphomas has been found to inhibit Kv1.3 channels, leading to lymphoma cell apoptosis (Wang et al., 2012). Currently, rituximab is also being used as a treatment for MS. Although the impact of rituximab on Kv1.3 regarding MS remains unexplored, existing research indicates that it minimises inflammatory activity, relapse frequency and CNS lesions (Chisari et al., 2022). This body of evidence underscores the significance of additional research on B lymphocytes in MS, while also establishing the patterns of expression and function for Kv channels within these subtypes of cells.

The electrophysiology study presented significant challenges, particularly due to the non-adherence of these cell subtypes which made it difficult to obtain additional data. Furthermore, it is crucial to consider variability when conducting human sample-based studies. The inherent diversity among individuals, including genetic, physiological, and environmental factors, can greatly influence research outcomes and complicate the interpretation of results (Bennett, 2010). Hence, we posit that the challenges in conducting experiments and the human variability involved may have contributed to the absence of noteworthy outcomes for T lymphocytes. Moreover, analysing T lymphocytes without considering their distinct cellular subtypes and activation states could also be a contributing factor. Previous studies on T lymphocyte electrophysiology have only examined Kv1.3 currents in small resting T lymphocytes (without regard for lineage, including naïve and memory T lymphocytes), resulting in an observed increase of current density in MS patients correlating with elevated Kv1.3 mRNA expression (Markakis et al., 2021). Conversely, Varga et al. (2009) analysis of Kv1.3 currents demonstrates lower levels in regulatory T lymphocytes compared to those found in naïve T lymphocytes from MS patients, whereas in healthy controls regulatory T lymphocytes and naïve T lymphocytes had comparable currents (Varga et al., 2009).

The expression of Kv channels in PBMCs is not significantly different based on clinical variables, MS types, or different therapies used. Our study acknowledges limitations such as variability inherent to human samples and challenges associated with electrophysiological recordings. Futures research should address these limitations by increasing samples size, investigating Kv channel expression and function within specific T lymphocyte subtypes and activation states, and exploring the role of Kv channels in B lymphocyte differentiation, proliferation, and cytokine production to shed light on their contribution to MS.

As a concluding remark, this study unveils novel insights into the altered Kv channel landscape in PBMCs and lymphocytes from MS patients. By highlighting upregulation of six Kv channel isoforms, sex-related differences, and the distinct electrophysiological profiles of CD3+ and CD19+ cells, our findings pave the way for further exploration of Kv channels as potential therapeutic targets in MS, particularly within specific immune cell subsets and in conjunction with sex-specific considerations.

## Acknowledgements

This work is part of the doctoral thesis of Marta Iglesias Martínez-Almeida, entitled “Characterization and study of peripheral proteins in schizophrenia lymphocytes”, presented at the University of Vigo in 2024.

## List of abbreviations

BBB: Blood brain barrier
BSA: Bovine serum albumin
CNS: Central nervous system
DAPI: 4,6-diamidino-2-phenilindol
EDSS: Expanded Disability Status Scale
IQR: Interquartile range
K_2_EDTA: Dipotassium ethylenediaminetetraacetic acid
Kv: Voltage-gated potassium
MS: Multiple sclerosis
NLR: neutrophil-lymphocyte ratio
PBMCs: Separate peripheral blood mononuclear cells
PVDF: polyvinylidene fluoride
TBS: Tris-buffered saline
TBST: Tris-buffered saline Tween

